# A Machine Learning Approach Predicts Tissue-Specific Drug Adverse Events

**DOI:** 10.1101/288332

**Authors:** Neel S. Madhukar, Kaitlyn Gayvert, Coryandar Gilvary, Olivier Elemento

**Author notes:** co-first authors (Neel S. Madhukar, Kaitlyn Gayvert). Correspondence: Olivier Elemento.

## Abstract

One of the main causes for failure in the drug development pipeline or withdrawal post approval is the unexpected occurrence of severe drug adverse events. Even though such events should be detected by in vitro, in vivo, and human trials, they continue to unexpectedly arise at different stages of drug development causing costly clinical trial failures and market withdrawal. Inspired by the “moneyball” approach used in baseball to integrate diverse features to predict player success, we hypothesized that a similar approach could leverage existing adverse event and tissue-specific toxicity data to learn how to predict adverse events. We introduce MAESTER, a data-driven machine learning approach that integrates information on a compound’s structure, targets, and phenotypic effects with tissue-wide genomic profiling and our toxic target database to predict the probability of a compound presenting with different types of tissue-specific adverse events. When tested on 6 different types of adverse events MAESTER maintains a high accuracy, sensitivity, and specificity across both the training data and new test sets. Additionally, MAESTER scores could flag a number of drugs that were approved, but later withdrawn due to unknown adverse events – highlighting its potential to identify events missed by traditional methods. MAESTER can also be used to identify toxic targets for each tissue type. Overall MAESTER provides a broadly applicable framework to identify toxic targets and predict specific adverse events and can accelerate the drug development pipeline and drive the design of new safer compounds.

## INTRODUCTION

Drug adverse events are currently one of the main causes of failure in drug development and are one of the top 10 causes of death in the developed world^1, 2^. Toxicity issues remain a leading cause for the rising clinical trial attrition rates^3, 4^. Even after a drug has been approved, adverse drug reactions remain a large burden on the medical system with the costs amounting to as much as $30 billion dollars annually in the USA^5^. Furthermore the identification of the serious adverse events associated with drugs frequently does not occur until after FDA approval, with as many as 50% of adverse events going undetected during human trials^6^. Due to the prevalence and impact of this problem, the U.S. Food and Drug Administration (FDA) has established the US FDA Adverse Event Reporting System (FAERS).

Most adverse event detection experiments are carried out in pre-clinical phases based on animal results or during early clinical trials. However not all adverse events are detected, due to several factors including limited relevance of animal models to human physiology, limited sample sizes during trials, and patient populations that may not be representative of the overall population^5^. Further complications may include the low frequency or late onset of some adverse events^5^. As a result, retrospective studies are currently an important method for further characterization of the side effects associated with drugs. However this requires a large number of patients to be treated first and is dependent on voluntary reporting, which is especially problematic as only 10% of all drug adverse events are reported post-approval^7^.

Ideally possible adverse events would be detected during the pre-clinical phases of drug development, even before animal studies. Cell lines and reporter assays may help detect unwanted side effects early, but are often imprecise. Computational screening methods are also critical components of current drug development pipelines for evaluating pre-clinical toxicity. In particular, drug-likeness measures, which use molecular features to estimate oral bioavailability as a proxy for drug toxicity, have been widely adopted. Examples of drug-likeness methods include Lipinski’s Rule of Five^8^ and the Quantitative Estimate for Drug Likeness^9^. More recently machine learning based methods have been proposed for predicting drug toxicity, including previous work from our group (PrOCTOR) which integrates established molecular properties with target-based features to directly predict broad clinical trial toxicity^10^. Other groups have developed diverse methods focused on predicting toxicity specific to the liver^11^. However no method has yet been developed with the granularity to predict multiple specific adverse events across different tissue types, such as heart attacks or neutropenia, for a specific drug. Better methods for predicting such adverse events could improve fast-fail procedures and facilitate better trial design. To address this problem, we introduce MAESTER, a new machine-learning platform for the prediction of tissue-specific drug adverse events. We show that for a set of 6 serious adverse events MAESTER achieves unprecedented accuracy while maintaining high specificity and sensitivity. Additionally we demonstrate how MAESTER could have identified drug adverse events that were missed by traditional screening methodologies but led to costly market withdrawal.

## RESULTS

### Identifying determinants of tissue-specific toxicities and adverse events

We first sought to identify drugs or compounds that are specifically toxic within individual tissues and compare them with compounds with no reported toxicities in these tissues. We focused on a set of six tissues whose corresponding AEs are correlated with clinical trial failures: liver, kidney, blood, heart, lung, and pancreas (**Fig.S1A**). We used the SIDER database of drug side effects to identify subsets of drugs that are associated with tissue-specific adverse events (TSAEs) (Table 1)^12^. For example we identified all drugs that have been associated with liver toxicities. For each tissue, we also established a “safe” set of drugs for comparisons identifying any drugs not associated with those TSAEs or other AEs highly correlated with fatalities in openFDA (https://open.fda.gov/) defined as having a fatality frequency > 13% (Fig.1A). For each drug, we compiled structural representations in the format of SMILES from DrugBank, differential gene expression profiles from the Broad Institute’s Connectivity Map (CMAP)^13^, growth inhibition patterns across the NCI60 cell lines (NCI60) from the NCI’s Developmental Therapeutics Program^14^, and bioassay data from PubChem^15^.

**Figure 1.**
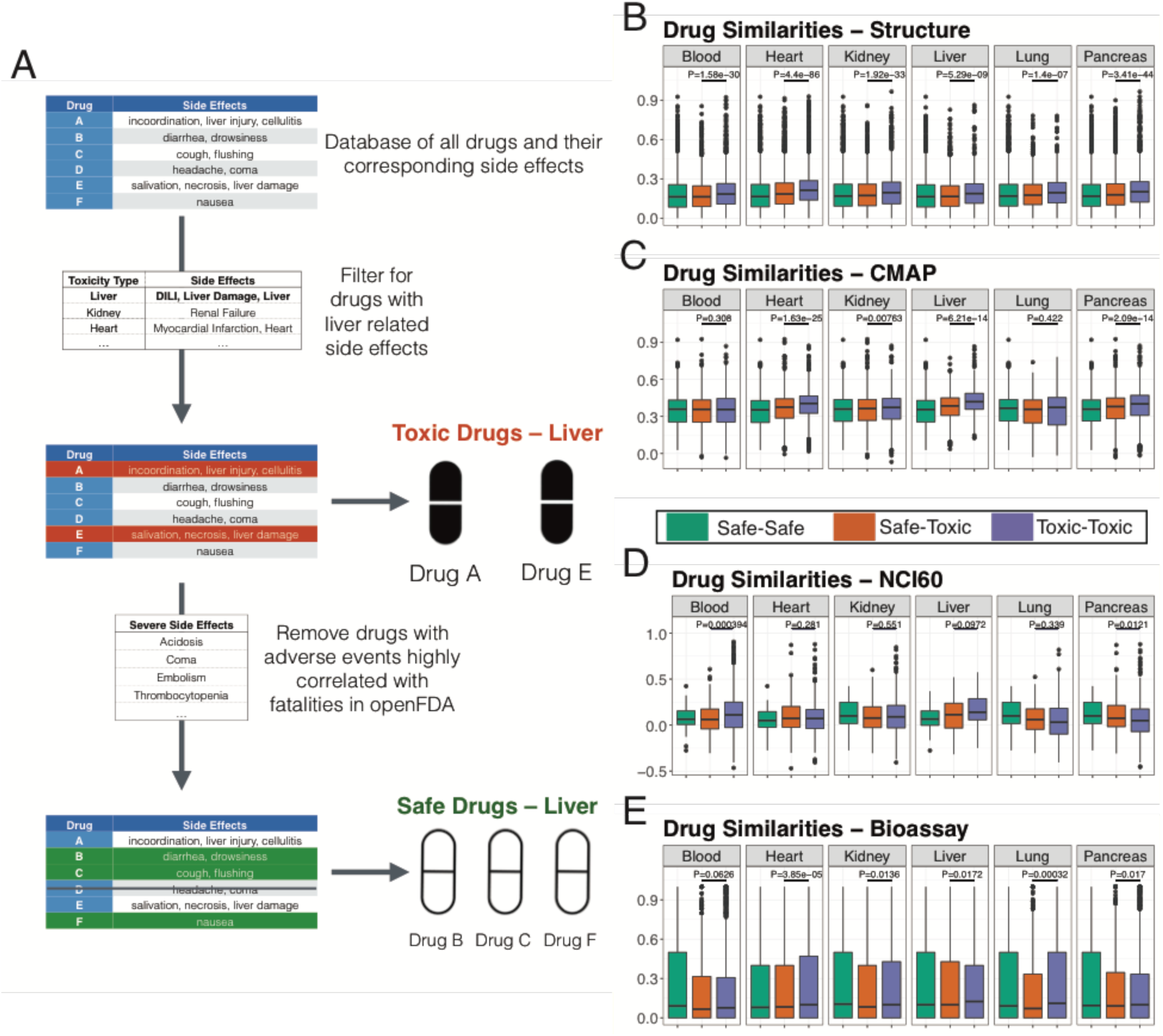
A) Schematic describing the process by which we selected our toxic and safe drugs for each specific tissue. B) Similarities of across all toxic drugs pairs, safe drug pairs, and all combinations of toxic and safe drugs for drug structures, C) gene expression changes, D) growth efficacies, and E) bioassays. P values were calculated using a Wilcoxon Rank Sum test.

**Table 1.**
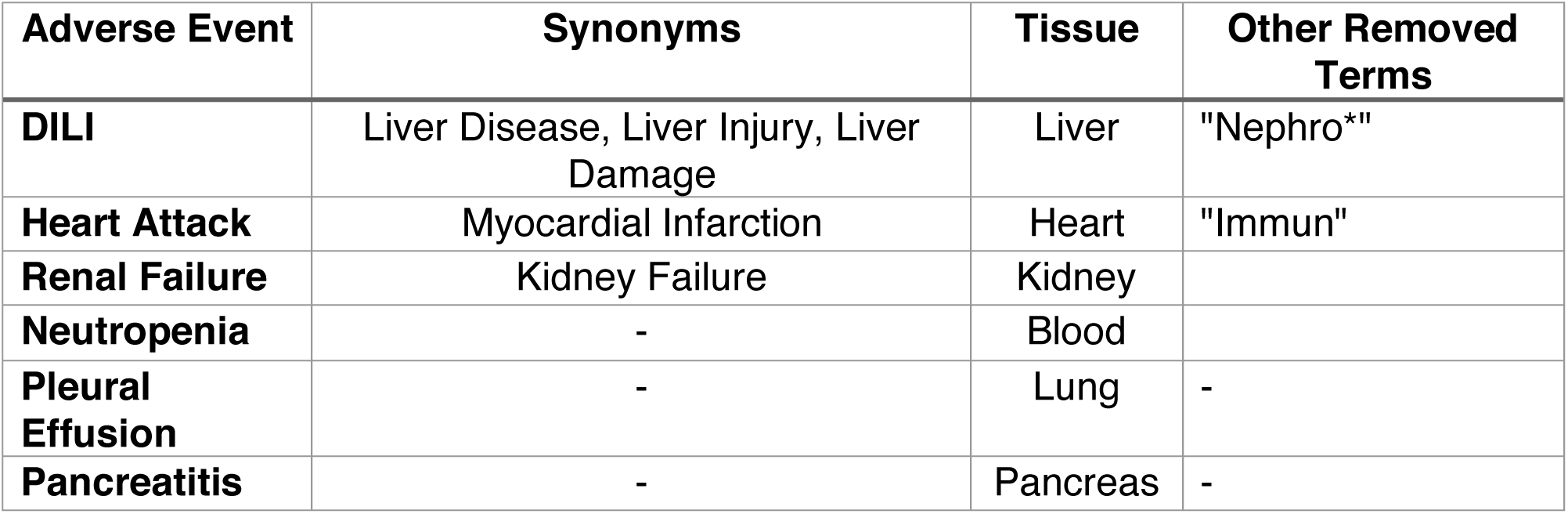
MAESTER Training Set Definitions. Table of the 6 major adverse event categories. In addition to the given adverse event, certain synonymous adverse events were also included and any drugs with containing an adverse event in the “other removed terms” category were removed excluded from the safe set.

For each tissue we then investigated how these safe and toxic drugs compare to each other. For each pair of drugs, we calculated a similarity score for each of the considered data types (**Methods**). We found that in all tissues, tissue-specific toxic drugs were most structurally similar to each other (Fig.1B). Additionally, toxic drugs tended to also be most similar to other toxic drugs in terms of differential gene expression profiles (Fig.1C), growth inhibition screens (Fig.1D) and bioassays (Fig.1E). Interestingly we found distinct patterns across the different tissue types – for instance, growth inhibition was best able to separate out drugs with blood specific adverse events, whereas gene expression changes had the greatest utility in the liver. These patterns could be incredibly valuable for adverse event prediction as they highlight how we can model the diversity across drugs with a given side effect. For example high structural similarity between a new compound and compounds known to be toxic in the heart could indicate potential cardiac toxicity for that new compound. Additionally high similarity between the compound-induced expression changes of a new compound with expression changes of compounds with known liver toxicity could suggest liver toxicity for the new compound.

We next examined how expression of a drug’s targets could be used to predict TSAEs. For this analysis we integrated tissue-specific expression data measured by the GTEX database. For each toxic or safe drug in a given tissue set (Fig.1A), we quantified the expression of all of that drug’s targets in the specific tissue. Overall drugs with adverse events in a specific tissue tended to also have higher target expression in that tissue than their safe drug counterparts (Fig.2A-E). This information helps illustrate how it is important to consider target based features and tissue-specific expression when predicting adverse events. This analysis also confirms that high expression of a drug’s target in a given tissue can help predict toxicity in that tissue.

**Figure 2.**
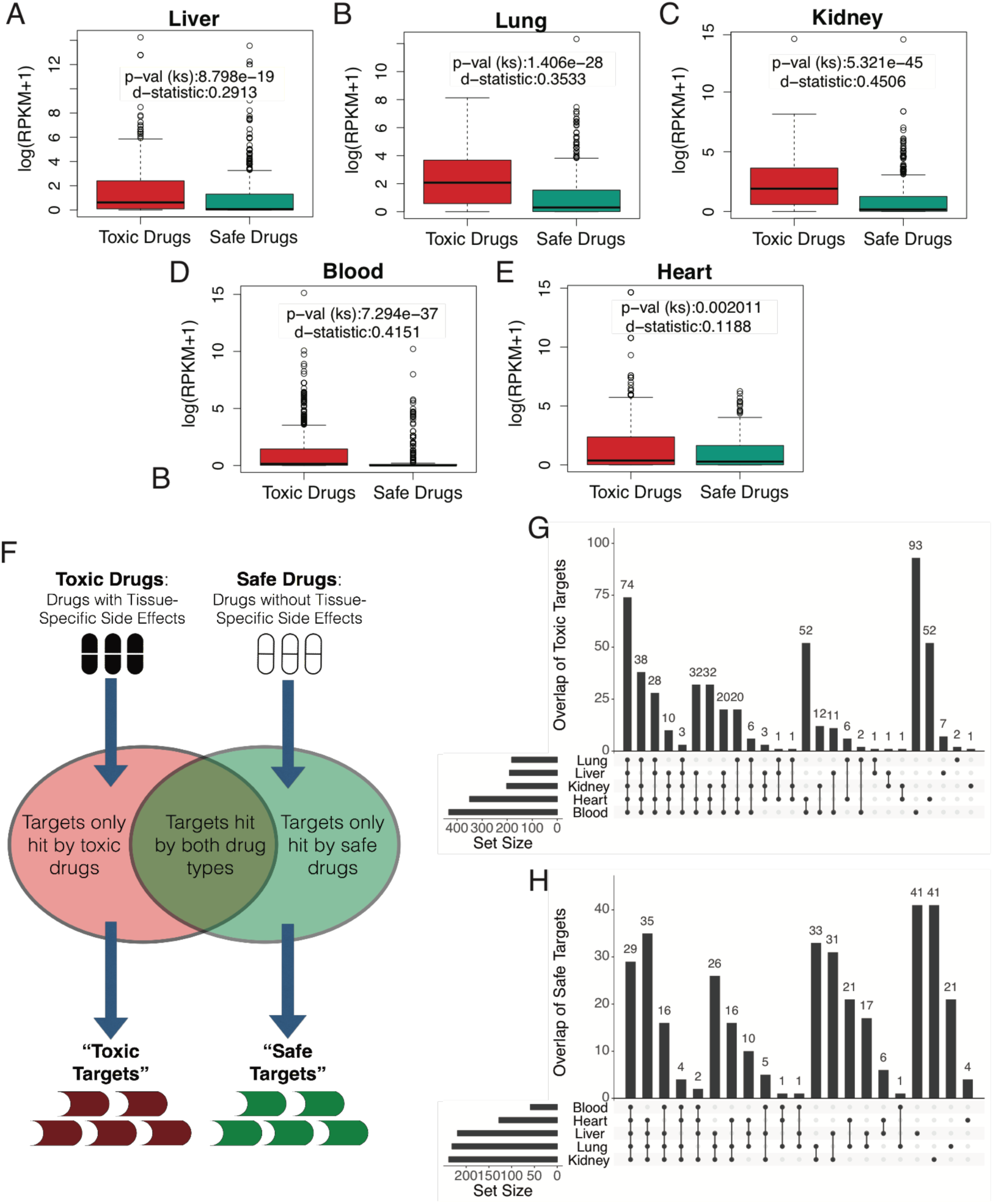
A–E) Distribution of target expression in a specific tissue for drugs with and without any tissue-specific adverse events (in that given tissue). P values and D statistic calculated using a KS test. F) Schematic for the selection of toxic and safe targets. G)UpSetR plot highlighting the overlap across tissue types for their respective toxic and H) safe targets.

### Distinct Patterns of Tissue-Specific Toxic and Safe Target Sets

Due to the significant relationship between drug target expression and related tissue adverse events, we next sought to define a set of tissue-specific “toxic targets”– proteins that are only targeted by drugs with known toxicity in that tissue – and “safe targets” – proteins only targeted by drugs with no related tissue toxicities. To do this, we begin by taking the safe and toxic drug sets described in Fig.1A and identifying any targets exclusive to each drug subset (Fig.2F). Interestingly we found that though there was a significant degree of overlap between the toxic and safe gene sets across multiple tissues, there were a number of proteins identified that were specifically associated with toxicity or non-toxicity in a single tissue (Fig.2G-H). For instance, ABL1 was flagged as a toxic target in all six tissues, whereas KCNJ3 and KCNJ6 – proteins involved in voltage gated potassium channels and the regulation of heartbeats – were only marked as toxic targets in the heart.

To further investigate features of tissue-specific toxic targets we expanded the procedure described in Fig.2F to generate toxic and safe targets for 30 different tissue types – including the 6 prior tested tissues. For each target, we computed a number of features, including tissue-specific expression, network properties (betweenness and degree), loss of function (LoF) mutation frequency, and essentiality status. We found that toxic gene sets tend to be more connected in an aggregated protein-protein interaction network (Fig.3A-B), be more intolerant for LoF mutations (Fig.3C), and be enriched for essential genes (Fig.3D). Finally, we used the ConsensusPathDB framework^16^ to measure for GO term enrichment and observed that for toxic gene sets the most commonly enriched terms had to due with cell death, receptor signaling, and apoptotic processes (Fig.3E) – pathways one would expect to be related to toxicity – whereas safe targets did not appear to be related to any toxicity related processes (Fig.3F) – likely due to the diverse nature and function of safe targets. Altogether these results suggest that tissue-specific toxic targets have specific recognizable features and that such features may be used to predict whether a new compound whose targets are known is likely to be toxic in a given tissue.

**Figure 3.**
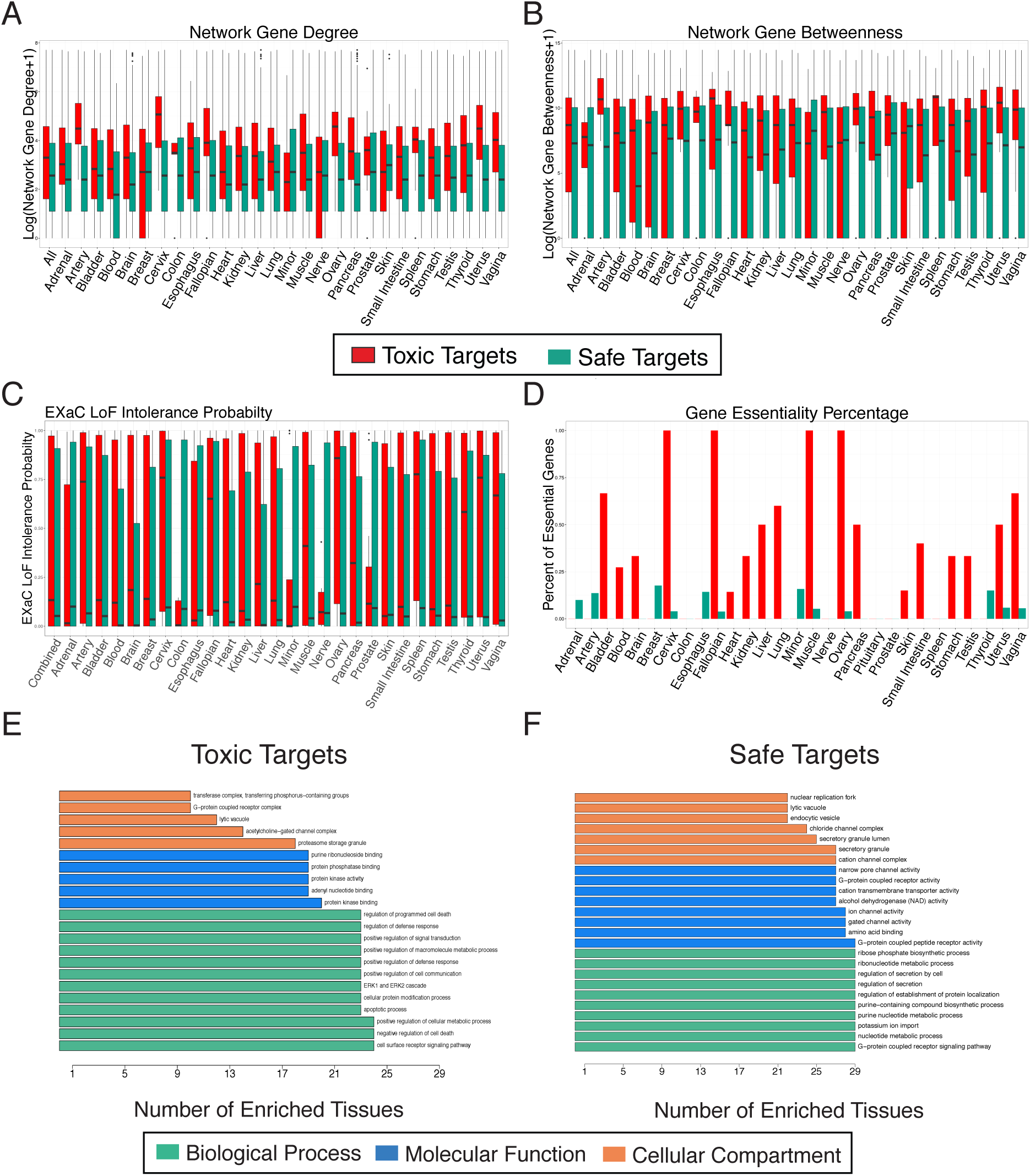
A-D) Distribution of features across multiple tissues for their individual toxic and safe targets. E) Number of tissues whose respective toxic or F) safe targets are enriched for a specific Gene Ontology category.

### Computational approach predicts likelihood of specific adverse events

To utilize these findings and more directly address the problem of adverse event prediction, we developed MAESTER (a **M**oneyball **A**pproach for **E**stimating **S**pecific **T**issue adverse **E**vents using **R**andom forests) to compute the probability of a compound presenting with a specific adverse event (Fig.4A). To do this, we expanded upon the framework of our previously published work on predicting broad clinical trial toxicities, PrOCTOR ^10^, and narrowed down the classification task to a set of specific adverse events that are correlated with clinical toxicity and have high reported frequencies of fatality in openFDA: drug-induced liver injury (DILI), nephrotoxicity, neutropenia, heart attack, pleural effusion, and pancreatitis (**Fig.S1A**). We began by using the framework described in Fig.1A to define a training set of safe and toxic drugs for each adverse event and its corresponding tissue. For the toxic drugs, we directly queried the database for drugs that are linked to each adverse event or its synonyms. We then took drugs that are not associated with any adverse event in the related tissue or any other severe adverse events to be the set of safe drugs (**Fig.S1B**). The set of keywords used to construct these training sets are described in Table 1.

**Figure 4.**
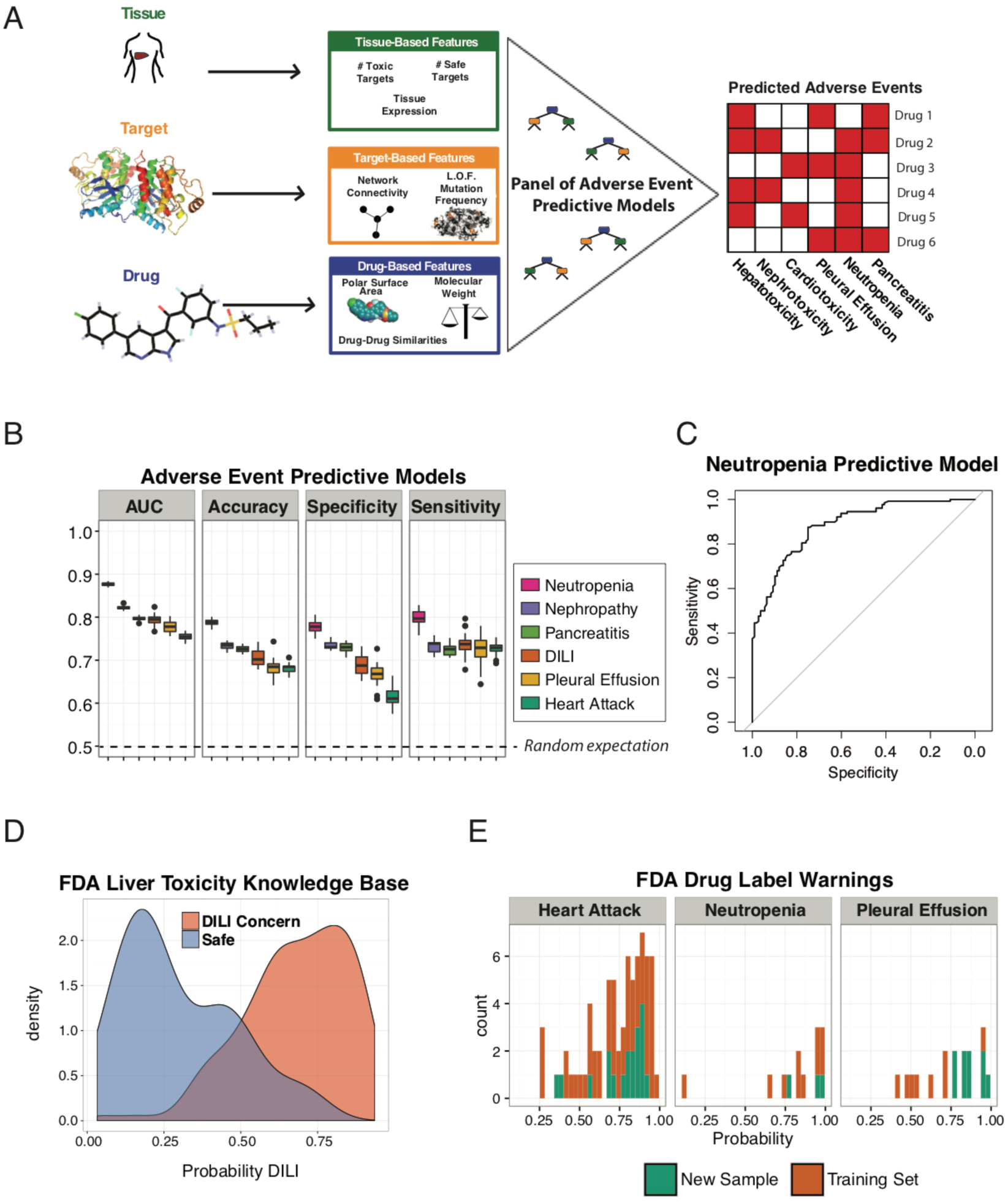
A) Schematic of MAESTER’s method of integrating multiple feature types to predict tissue-specific adverse events. B) Performance metrics for multiple MAESTER prediction models. C) Area under the receiver-operating curve for MAESTER’s Neutropenia model. D) Distribution of MAESTER DILI probabilities for drugs marked as “DILI Concern” or “Safe” by the FDA Liver Toxicity Knowledge base. E) MAESTER Predictions for drugs with FDA warning labels for heart attacks, neutropenia, or pleural effusion.

Building upon the framework of PrOCTOR, MAESTER integrates 13 structural features, 35 target and tissue features, and 8 drug similarity properties to produce a suite of classifiers that are able to predict the likelihood of each adverse event (Fig.4A). Given the established validity of drug-likeness measures in capturing toxicity, we also included properties considered by the Lipinski^8^, Veber^17^, and Ghose^18^ rules, and the Quantitative estimate for Drug-Likeness (Q.E.D.)^9^ as well as the measures themselves. For tissue-based features, we considered the number of known drug targets that fall in the associated tissue-specific safe and toxic gene sets we created earlier. We also included the above described tissue expression features from GTEx^19^, network properties (connectivity and degree), and loss of function mutation frequency^20^. Finally we integrated the different similarity scores (structural, CMAP, NCI60, and bioassay) through two different measures. The first similarity metric represents whether the drug is more similar to known safe or toxic molecules by using a signed Kolmogorov-Smirnov D-statistic. The second similarity metric is a count of the number of highly similar drugs with known TSAEs.

The classifiers were evaluated using 10-fold cross validation. All adverse events achieved significant predictive performances with an average accuracy of 72% and area-under-the-receiver-operator curve (AUC) of .81 (Fig.4B, Table 2). Focusing specifically on neutropenia – a major cause of clinical trial failure and mortality in cancer and immunocompromised patients^21^– MAESTER achieved an AUC, accuracy, specificity, and sensitivity of 0.8843, 0.7839, 0.7778 and 0.7891 respectively (Fig.4C, Table 2) – to our knowledge the highest reported results for the computational prediction of neutropenia.

**Table 2.** MAESTER Model Performances. AUROC, Accuracy, Specificity, and Sensitivity values for each of MAESTER’s underlying models.

| **Adverse Event** | **# safe**<br>**drugs** | **# toxic**<br>**drugs** | **AUROC** | **Accuracy** | **Specificity** | **Sensitivity** |
| --- | --- | --- | --- | --- | --- | --- |
| **DILI** | 105 | 268 | 0.8079 | 0.7373 | 0.7333 | 0.7388 |
| **Heart Attack** | 113 | 166 | 0.7552 | 0.6882 | 0.6283 | 0.7289 |
| **Renal Failure** | 127 | 165 | 0.8198 | 0.7329 | 0.7087 | 0.7515 |
| **Neutropenia** | 108 | 128 | 0.8843 | 0.7839 | 0.7778 | 0.7891 |
| **Pleural**<br>**Effusion** | 128 | 59 | 0.7761 | 0.6631 | 0.6562 | 0.678 |
| **Pancreatitis** | 126 | 153 | 0.7967 | 0.7348 | 0.7302 | 0.7386 |

### MAESTER identifies adverse events in independent test sets

We further assessed MAESTER’s performance using an independent validation test set. For liver toxicity, the FDA has curated the Liver Toxicity Knowledge Base (LTKB) that classifies a number of compounds based on their risk of causing liver toxicity. We found that MAESTER can significantly distinguish drugs that are of DILI-concern from those classified as no concern using this independent database (Fig.4D) (*p* < 2.2e-16, Mann-Whitney U test). For heart attacks, pleural effusion, and neutropenia we turned to FDA drug label warnings as reported in openFDA. We found that MAESTER correctly identified 76.3% of drugs with heart attack risk (*p*=0.04589, Binomial test), 75.0% with pleural effusion risk (*p*=0.01474, Binomial test), and 87.5% with neutropenia risk (*p*=0.0782, Binomial test) (Fig.4E). These tested compounds did not have their specific adverse events listed in SIDER and thus were not in our original training set, further highlighting MAESTER’s potential to predict adverse events for new compounds.

A feature importance analysis revealed that there is a subset of features that were consistently predictive across all of MAESTER’s adverse event models (**Fig.S2A**). The toxic and safe gene sets, structural and bioassay similarity features, polar surface area, and expression of the drug target in mature B cells are important in a majority of models. We also identified a subset of features that are uniquely predictive in specific models. For example, target expression in digestive organs (e.g., colon, small intestine, stomach) were highly important in the prediction of DILI (**Fig.S2B**), expression in immune-related cells (centroblasts, T cells, spleen) were important for neutropenia prediction (**Fig.S2C**), and the network degree of the drug target was the most important feature in prediction of pleural effusion (**Fig.S2D**).

We then compared the predictions for drugs across all models (**Fig.S3A**). We found that there were subsets of drugs that are predicted to be safe or toxic by most or all models. We found that drugs predicted to have many TSAEs tended to have higher predicted toxicity levels (measured by the PrOCTOR score) (**Fig.S3B**) than drugs that were predicted to have one or less TSAEs (**Fig.S3C**, *p*=1.178e-06, Mann-Whitney U test).

### MAESTER predicts specific adverse events for withdrawn drugs

To test MAESTER’s ability to detect adverse events that may have been missed by traditional approaches, we next focused on drugs that been approved but were later withdrawn due to toxicity concerns. This is especially relevant because cardiotoxicity and hepatotoxicity – two of MAESTER’s adverse event models – are the largest causes of toxicity related withdrawal ^22^. We began by focusing on two well-known cases of drug withdrawal – Vioxx and Avandia, both withdrawn for cardiac toxicity– and found that MAESTER scored each as highly likely to cause cardiac toxicity (Fig.5A-B). In fact, comparing Avandia (Rosiglitazone) to a less toxic analog (Pioglitazone) we observed that the difference in reported toxicities corresponded to a difference in their MAESTER scores. We found that these predictions did not change substantially when we removed both drugs (and their analogs) from the original training set, retrained MAESTER’s underlying model, and rescored each compound. To further expand this analysis we curated a list of withdrawn drugs (that were not part of MAESTER’s original training set) and their reason for withdrawal (**Methods**). For each drug we computed a MAESTER probability corresponding to the specific reason for withdrawal (Table 3). We found that for 87.5% of the withdrawn drugs MAESTER predicted that specific adverse event with a probability greater than 0.5 – significantly more than would have been expected by random chance (*p* =0.0003, Fisher’s exact test). To further evaluate MAESTER’s ability to flag withdrawn drugs, we compared MAESTER probabilities of withdrawn drugs against probabilities for drugs of similar indications that were never withdrawn and were not known to have the reported adverse event (Fig.5C-F). We found that withdrawn drugs had significantly higher MAESTER adverse event probabilities than approved drugs of the same indication (*p* =0.0027 and 0.0424, Fisher’s exact test). Overall these results highlight MAESTER’s ability to specifically identify compounds with adverse events that were missed by traditional approaches.

**Figure 5.**
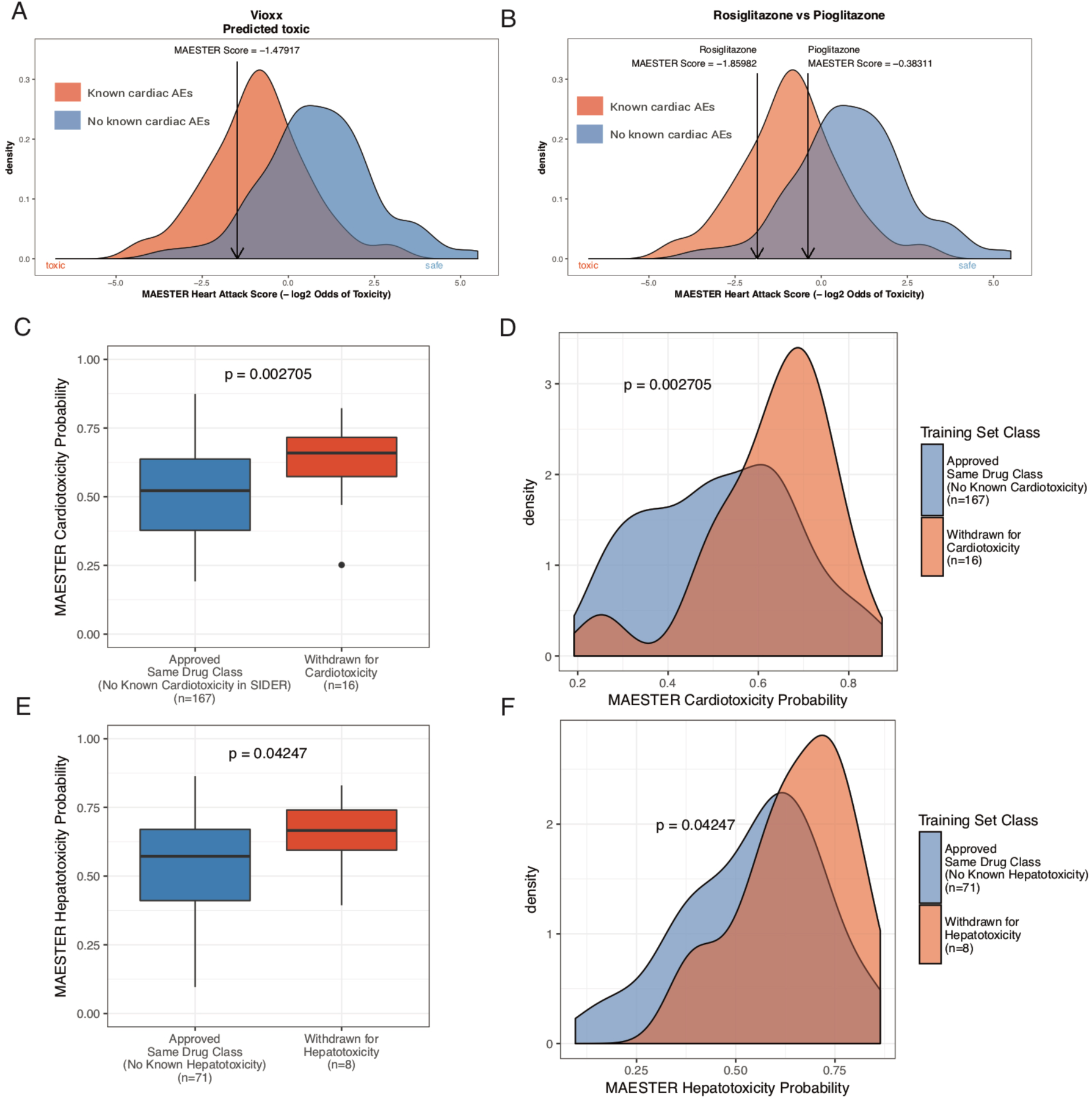
A) Distributions of MAESTER scores for all drugs known to cause heart attacks and those considered safe. MAESTER scores for Vioxx, B) Rosiglitazone, and Pioglitazone are indicated with arrows. C-D) MAESTER scores for drugs withdrawn for cardiac toxicity compared to approved drugs of the same class with no known cardiac toxicities. E-F) MAESTER scores for drugs withdrawn for liver toxicity compared to approved drugs of the same class with no known liver toxicities. All p values were calculated using a Wilcoxon rank sum test.

**Table 3.**
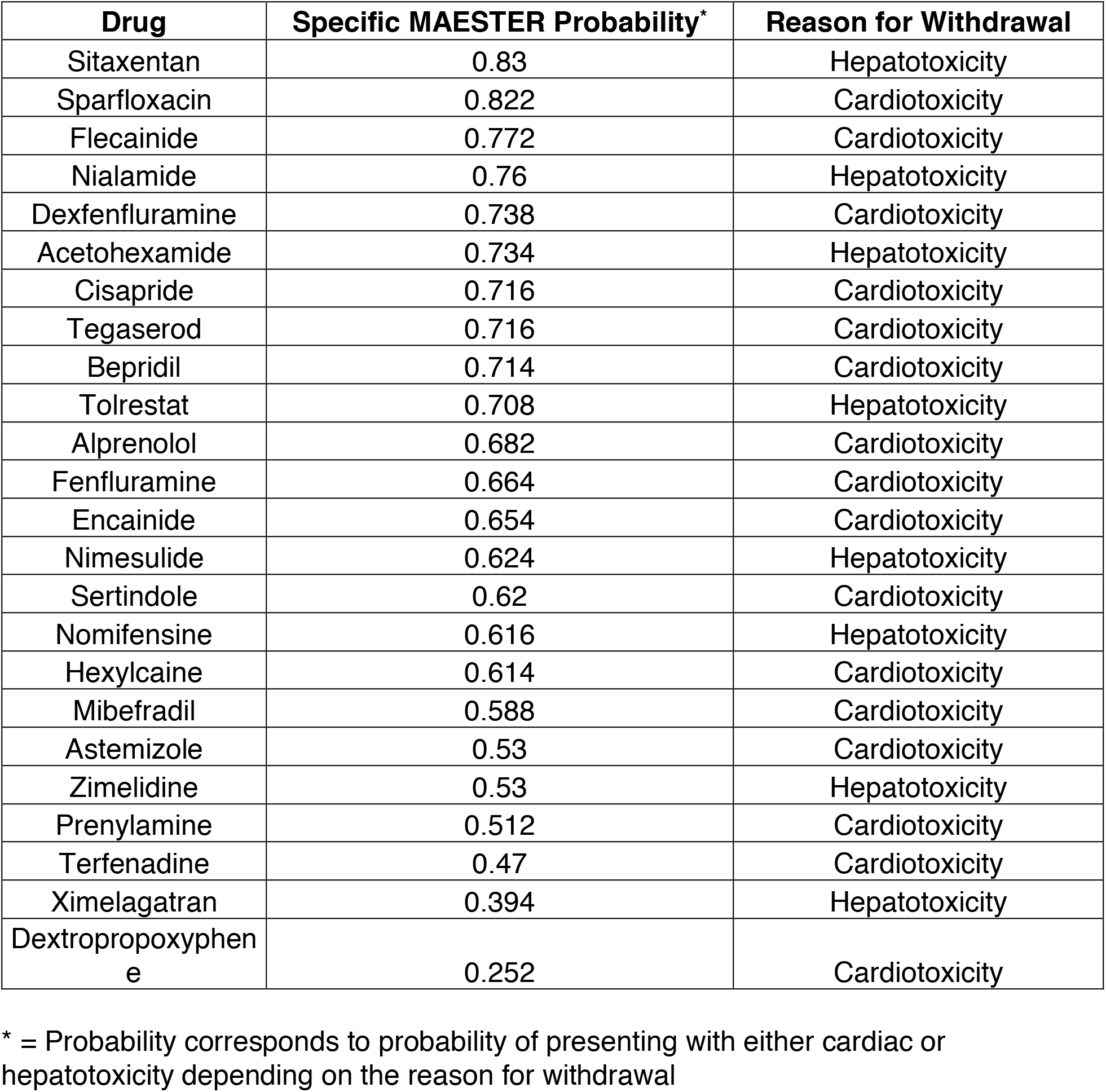
MAESTER Performance on Withdrawn Drugs. List of withdrawn drugs, their reason for withdrawal, and the corresponding MAESTER score.

## DISCUSSION

Pre-clinical toxicity screening is one of the most important parts of drug development. Existing experimental methods are cumbersome and often do not translate to clinical results. Computational methods for predicting toxicity can complement and perhaps guide experimentation to evaluate toxicities. However prior methods have for the most part focused only on molecular properties and predicting broad clinical toxicities rather than specific adverse events. We have proposed MAESTER, a data-driven machine learning approach that integrates information on a compound’s structure, targets, and downstream effects to predict the probability of a compound presenting with different adverse events. When trained on drugs with known adverse events, MAESTER performs at high accuracy, sensitivity, and specificity across six different prediction tasks. Additionally MAESTER performs with high accuracy on external FDA test sets and drug warning labels, and could accurately identify adverse events for withdrawn drugs that may have been missed during traditional analyses.

We have identified sets of toxic and safe drugs and genes that are associated with adverse events in specific tissues. We found that tissue-specific toxic drugs tend to be more similar to each other than known safe drugs and that their associated targets are more highly expressed in corresponding tissues. We found tissue-specific toxic targets tend to be enriched for apoptosis and cell death related biological processes, more connected in protein-protein interaction networks, and are classified as more essential. Leveraging this data, we developed MAESTER to combine compound and target properties to predict the likelihood of specific adverse events. Because it is trained on drugs with known adverse events, MAESTER can directly predict clinical effects compared to cell or animal screening methods whose toxicity predictions may not translate to the clinic.

One of the strengths of our big data approach is that it can consider a large number of features without prior bias. This will become especially powerful in the coming years as more large pharmacogenomics datasets become available to integrate. Analysis of these features can aid in future drug design by providing insight into what types of drugs are likely to be toxic and feeding this information back to the chemists. Additionally, while toxicity is often modeled as a broad feature, often times it is a patient specific effect. As more patient specific data becomes available MAESTER can be improved to predict patient specific adverse events. This could be used to guide clinical trial design by specifically selecting patients unlikely to present with toxic effects and radically change how people approach precision medicine.

