## Supplementary Materials for "A Machine Learning Approach Predicts Tissue-Specific Drug Adverse Events"

**SUPPLEMENTAL FIGURES**

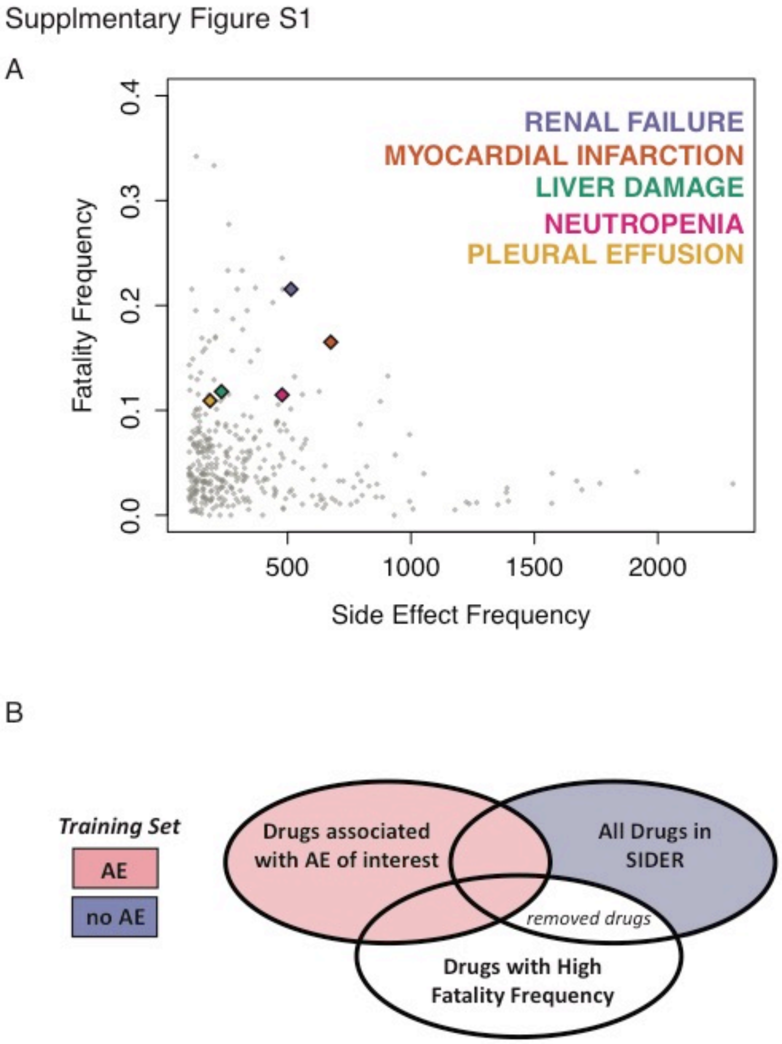


**Figure S1 –** A) Frequency of various side effects vs. frequency of fatalities in openFDA. B) Schematic describing selection of drugs for training set categories.

Supplementary Figure S2

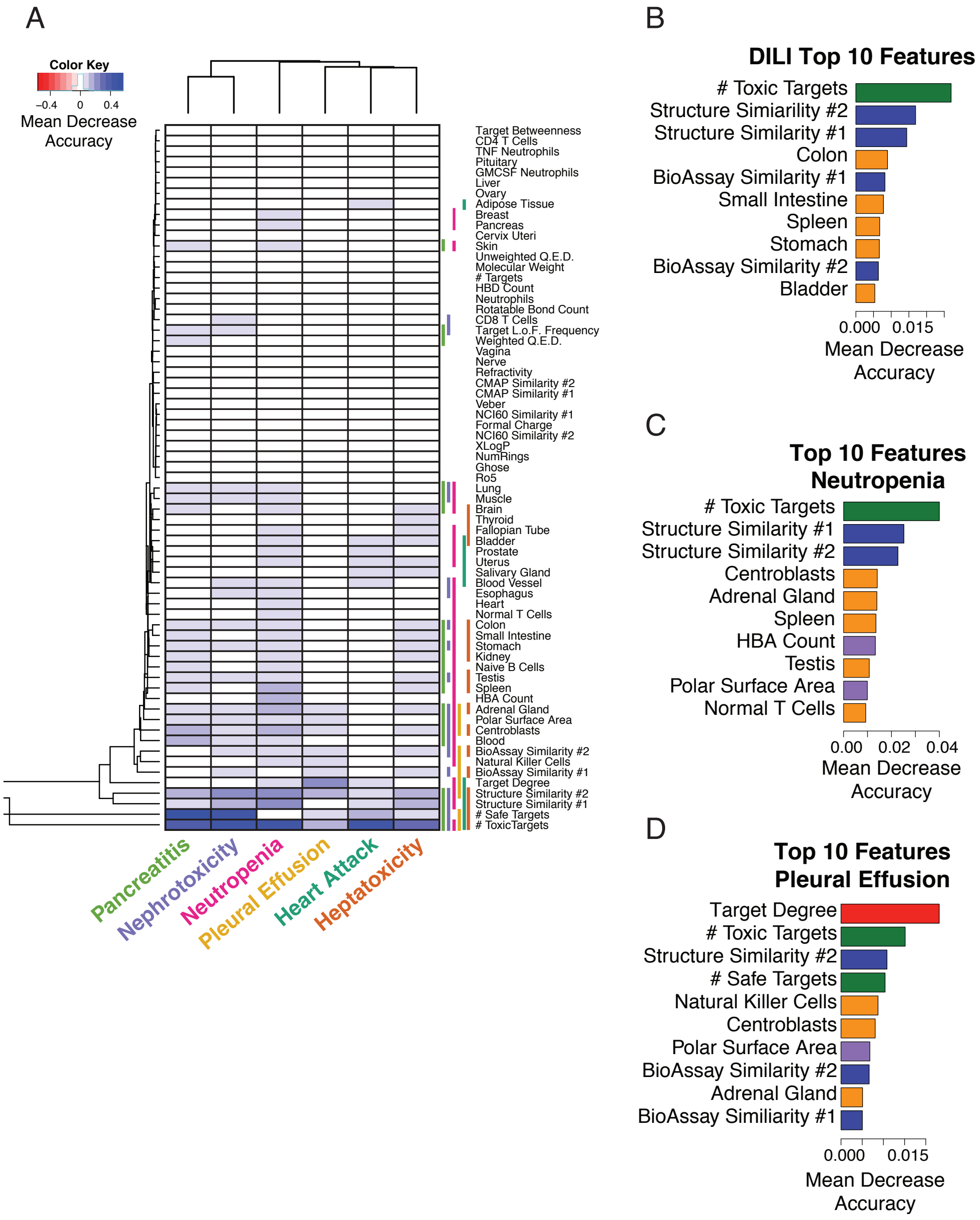


**Figure S2 –** A) Feature importance analysis for MAESTER's models and top 10 features for B) DILI, C) Neutropenia, and D) Pleural Effusion prediction models.

Supplementary Figure S3

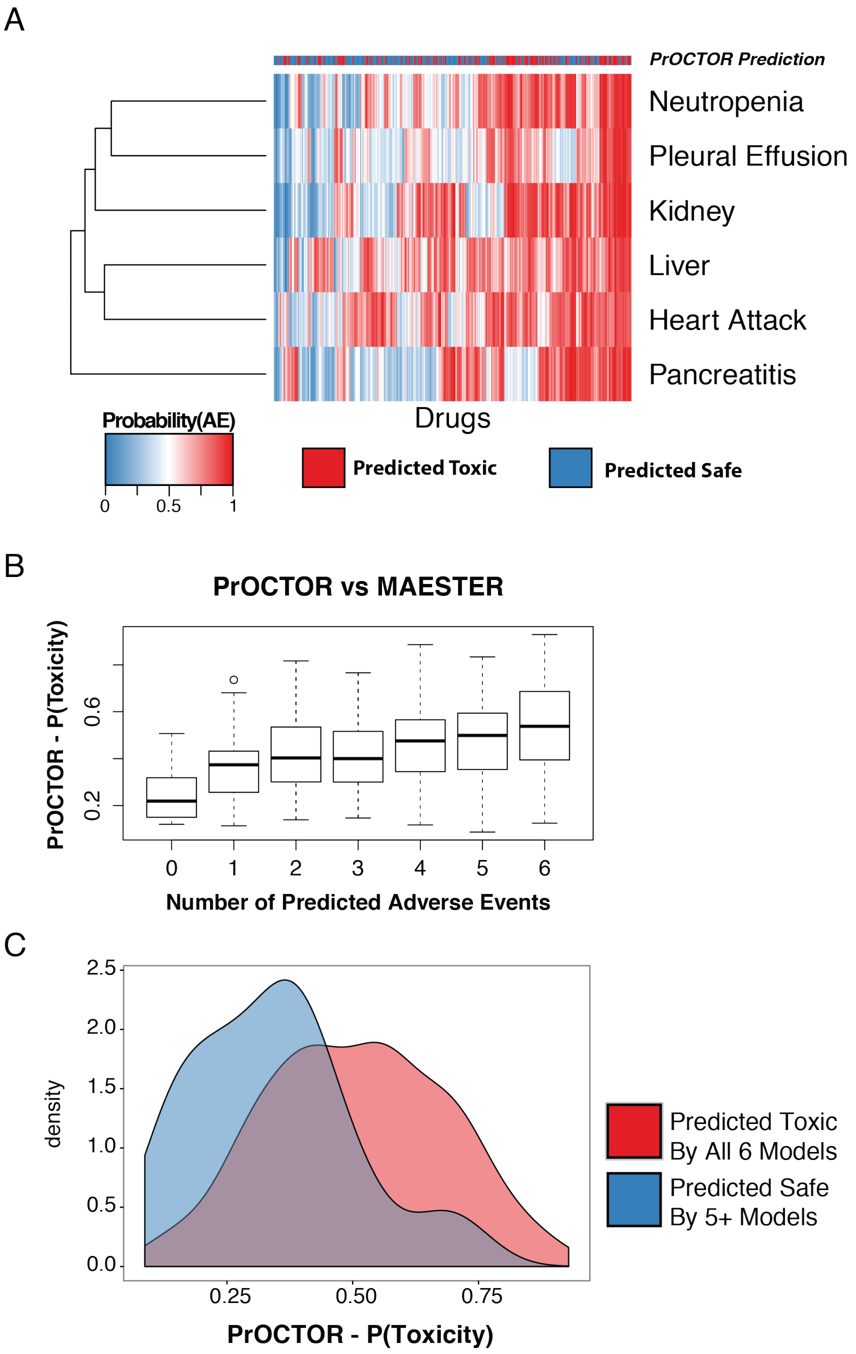


**Figure S3 –** A) MAESTER probabilities for all models across a number of drugs compared to the PrOCTOR prediction score. B) Distribution of PrOCTOR scores for drugs that predicted to 0-6 adverse events by MAESTER. C) Difference in PrOCTOR score distributions for drugs predicted to have 6 AEs by MAESTER vs. drugs that were predicted to have up to (but not more than) 1 AE by MAESTER.

### MATERIALS AND METHODS

#### Training Set

We downloaded the Side Effect Resource (SIDER) database from sideeffects.embl.de. The *meddra_adverse_effects.txt* table was used to extract reported adverse events and the *MedDRA Preferred Term* descriptor to group similar side effects. For each of the 30 major tissue types reported in the Genotype-Tissue Expression (GTEx) project, we identified the drugs that are associated with any adverse events by searching for keywords associated with the tissue name. We used the openFDA resource (https://open.fda.gov/) to identify drugs in SIDER that are associated with high fatality rates, which we defined to be greater than 13% of reported cases. We further extracted the names of the drugs associated with specific adverse events, including synonyms listed in Table 1. To define a set of drugs not associated with a given adverse event, we took the remaining drugs in SIDER and removed those that are associated with any other toxicity in the tissue, according to terms listed in Table 1, or high fatality rates, as defined using the openFDA resource above.

#### Feature Derivation

##### *Chemical Features*

The structures (sdf format) were downloaded for all of the drugs in DrugBank. The molecular weight, polar surface area, hydrogen bond donor and acceptor counts, formal charge and number of rotatable bounds were extracted from the sdf file for each of these compounds. When that information was missing, it was filled in using PubChem or by computationally estimating these values using ChemmineR in R. Drug-likeness rule outcomes for the Lipinski, Veber, and Ghose rules were derived using these features. The QED values were computed using the author-released script.

##### *Network features*

We constructed the aggregated biological network by taking the union across multiple databases of gene-gene interactions. [1-3]. The network degree of a gene was calculated as the number of gene neighbors that a particular gene has. The network betweenness for a particular gene (i.e. vertex) is defined as the number of shortest paths that travel through the vertex. For drug, we considered the maximum network degree and betweenness of its target genes. These measures were calculated using R's igraph package[4].

##### *Tissue features*

The Genotype-Tissue Expression (GTEx) project[5] dataset of 2921 RPKM RNA-Seq samples were downloaded from http://www.gtexportal.org/home/. For each tissue, the median RPKM was calculated for each gene. For each drug, the maximum RPKM of its targets was considered.

*Target Loss Frequency*

The Exome Aggregation Consortium (ExAC) database [6] was downloaded from www.**exac**.broadinstitute.org. The loss frequency was calculated to be percentage of deleterious mutations for each gene.

*Drug Similarities*

All drug pair similarities were calculated as described by Madhukar et al[7]:

1. Growth Inhibition Data: For each pair of drugs we calculated a pearson correlation value across the 60 data points (Figure 2.1).

2. Gene expression and Chemogenomic Fitness Scores: A pearson correlation was used to measure the degree of similarity for the profiles of two drugs.

3. Bioassays: All bioassays were classified as either positive or negative based on the data available in Pubchem. A jaccard index was calculated based on the number of shared "positive" assays between two drugs. We required that each drug pair have been tested in at least one similar assay for a similarity score to be calculated.

4. Chemical Structures: For each drug we extracted the isomeric SMILES and used the atom-pair method [8] to calculate the structural similarity between two compounds (Figure S1).

#### The MAESTER Approach

For each adverse event listed in Table 1, we trained a model using the training set and features described above. It was trained using the random forest model, an ensemble decision tree based approach, which constructs 50 bootstrapped decision trees. A sub-sampling approach was used to account for any imbalance in the ratio of toxic drugs to safe drugs, by randomly down-sampling the larger class of samples. To reduce the odds of poor representatives being sampled, this was repeated 30 times. The labels were assigned by taking the consensus across the set of bootstrapped trees and replicates. This approach also yields a probability for each test sample. This probability was used to calculate an odds score = $\log_2\left(\frac{P(\text{approval})}{P(\text{failure})}\right)$.

#### Independent Datasets

The FDA annotated drug-induced liver toxicity (DILI) dataset was downloaded from the FDA website at http://www.fda.gov/ScienceResearch/BioinformaticsTools/LiverToxicityKnowledgeBase/ucm226811.htm. The openFDA resource (https://open.fda.gov) was used to identify drugs that have warning labels for the relevant toxicity events. We extracted the set of drugs that have been withdrawn for known liver or cardiotoxicity reasons from DrugBank descriptions for withdrawn drugs and supplemented with information curated from https://en.wikipedia.org/wiki/List_of_withdrawn_drugs. These were compared to MAESTER predictions for all drugs of the same class that have not been previously annotated in SIDER for the given adverse event.

#### Statistical Analyses

We used area under the receiver operating characteristic (ROC) curve and 10-fold cross validation to evaluate the predictive power of our approach. For the analysis of predictions for the FDA drug warning dataset, we tested for enrichment of predictions using the binomial test. We tested for differences in predictions in the FDA DILI dataset and between classes for the withdrawn drug datasets using the unpaired Student's t-test. We tested the enrichment for probabilities above 0.5 for the withdrawn drugs by using a Fisher's exact test comparing the number of withdrawn drugs with a MAESTER probability above 0.5 against the number of approved drugs (with no known cardiac or hepatotoxicity) with a MAESTER probability above 0.5.
